# SCORPIUS improves trajectory inference and identifies novel modules in dendritic cell development

**DOI:** 10.1101/079509

**Authors:** Robrecht Cannoodt, Wouter Saelens, Dorine Sichien, Simon Tavernier, Sophie Janssens, Martin Guilliams, Bart Lambrecht, Katleen De Preter, Yvan Saeys

## Abstract

Recent advances in RNA sequencing enable the generation of genome-wide expression data at the single-cell level, opening up new avenues for transcriptomics and systems biology. A new application of single-cell whole-transcriptomics is the unbiased ordering of cells according to their progression along a dynamic process of interest. We introduce SCORPIUS, a method which can effectively reconstruct an ordering of individual cells without any prior information about the dynamic process. Comprehensive evaluation using ten scRNA-seq datasets shows that SCORPIUS consistently outperforms state-of-the-art techniques. We used SCORPIUS to generate novel hypotheses regarding dendritic cell development, which were subsequently validated *in vivo*. This work enables data-driven investigation and characterization of dynamic processes and lays the foundation for objective benchmarking of future trajectory inference methods.

## 2 Introduction

During the past three decades, flow cytometry and imaging techniques have been instrumental in profiling and characterizing single cells in a high-throughput manner. Recent advances in RNA sequencing now enable us to profile the whole transcriptome of individual cells, which allows studying rare cells [1, 2] or unravelling heterogeneous cell populations [1, 3, 4]. Single-cell RNA sequencing (scRNA-seq) has shed new lights on biology in many fields including microbiology, neurobiology, immunology and cancer research [5]. One domain which has benefited greatly from advancements in single-cell transcriptomics is the study of dynamic processes [6] including cell development and differentiation [7, 8, 9, 10], responses to stimuli [3, 11], and cyclic processes such as the cell cycle [12].

Dynamic processes are traditionally investigated by developing a time series model [13]. Time series data are typically obtained by observing gene expression levels of bulk populations of cells at multiple time points. Despite their utility, time series experiments are still associated with several technical and biological challenges such as time-resolution, cellular heterogeneity and the need for synchronization conditions. As a result, researchers now flock to computational methods which derive models of dynamic processes from single-cell data. By modelling a dynamic process as a trajectory and mapping the cells to regions in the trajectory, the progression of a cell in the dynamic process of interest can be predicted. Such computational methods, referred to as trajectory inference (TI) methods, can then be used to identify new marker genes associated with specific transition states [14], or novel intermediate states [8], and infer regulatory networks underlying the dynamic process [15].

Pioneering TI methods such as Monocle [16] and Wanderlust [16] have been instrumental in laying the foundations of the methodology, which typically consists of two main steps. In the dimensionality reduction step, the high-dimensional dataset (with thousands of genes) is converted to a low-dimensional representation using manifold learning techniques or graph-based techniques. In the subsequent trajectory modeling step, a model is constructed from the cells in the reduced space, by predicting the different cell states, inferring a trajectory through them, and projecting the cells on to the trajectory.

Wanderlust [16] requires a starting cell to be given as additional input. It creates a k-nearest-neighbor (KNN) graph to reduce the dimensionality, and orders cells according to their shortest-path distances to the starting cell. In order to improve the robustness of this approach, Wanderlust calculates a consensus ordering from the orderings obtained from bootstrapped KNN graphs. Monocle [17] uses Independent Component Analysis (ICA) to reduce the dimensionality. As the time complexity of ICA scales poorly with the number of genes in the dataset, Monocle first selects the genes most differentially expressed between given cell states. By calculating a minimum spanning tree between the cells and finding the longest connected path therein, the cells are ordered by projecting them onto the closest point in the path. Waterfall [18] reduces the dimensionality with Principal Component Analysis (PCA). Subsequently, the cells are clustered and a minimum spanning tree is calculated between the cluster centers. The longest path starting from the leftmost cluster is used as a trajectory, and the cells are ordered by perpendicularly projecting them onto the closest point on the trajectory.

Although these pioneering studies have shown that TI methods can be a powerful tool to improve our understanding of cellular dynamic processes [16, 17] the relative advantages and weaknesses of particular TI methods are still unclear at this point. In this study, we designed a benchmarking strategy for TI methods, which uses the known ordering of cellular states to evaluate the quality of the inferred ordering. When we used this strategy to assess the performance of state-of-the-art TI methods on a wide range of datasets, we found that none of the current methods performed well on all datasets consistently. We reasoned that there are two causes for this observation. First, at the time of development of these methods, the technologies that enabled profiling the transcriptome of single cells had just been released, and scRNA-seq datasets investigating dynamic processes were a scarce commodity. Existing TI methods were therefore evaluated on only one or two datasets, and might perform suboptimally on new datasets. Second, these methods use prior knowledge in order to obtain more robust models, while the methods were evaluated in an unsupervised setting. By providing the method with prior knowledge, bias towards existing knowledge is introduced into the model, which might preclude the researcher from discovering new information such as heterogeneities and hidden subpopulations in known cellular groupings.

## 3 Results

We introduce SCORPIUS, a novel method for inferring trajectories in a purely data-driven way, and we subsequently evaluate this method both computationally as well as biologically. To this end, we performed the first quantitative and extensive benchmark of TI methods on ten datasets. Subsequently, we demonstrate its practical usefulness by applying it to the dynamic process of dendritic cell development, and confirm the generated hypotheses *in vivo*. Compared to existing TI methods, SCORPIUS offers three main advantages. First, it produces accurate models in an extensive benchmark on a wide range of dynamic processes and predicts the progression of individual cells along those dynamic processes. Second, SCORPIUS does not require any user input and works in a purely data-driven fashion, which minimizes the amount of bias that may be introduced into the model and might lead to novel and unexpected findings. Finally, in order to improve the interpretability of the model, it is able to predict the involvement of genes in the dynamic process of interest, and visualize interesting gene expression patterns in an intuitive manner.

### 3.1 SCORPIUS constructs data-driven models of dynamic processes

Comparable to other TI methods, SCORPIUS assumes that a given dataset contains the genome-wide expression profiles of hundreds to thousands of cells, which were uniformly sampled from a linear dynamic process. Figure 1 presents the main steps of the SCORPIUS methodology: dimensionality reduction, trajectory modeling and gene prioritization. During the dimensionality reduction step (Figure 1b), the correlation distance between all pairs of cells is calculated. By default, SCORPIUS uses the spearman correlation as it is unit independent, and is typically more robust than other correlation distances when high levels of noise are contained within the dataset. Next, SCORPIUS removes outliers, as these could negatively impact the trajectory inference. Finally, multi-dimensional scaling (MDS) is used to reduce the dimensionality to n components. The reduced space highlights the main structure in the data and effciently reduces technical noise, making it easier to infer a trajectory in the next step.

**Figure 1:**
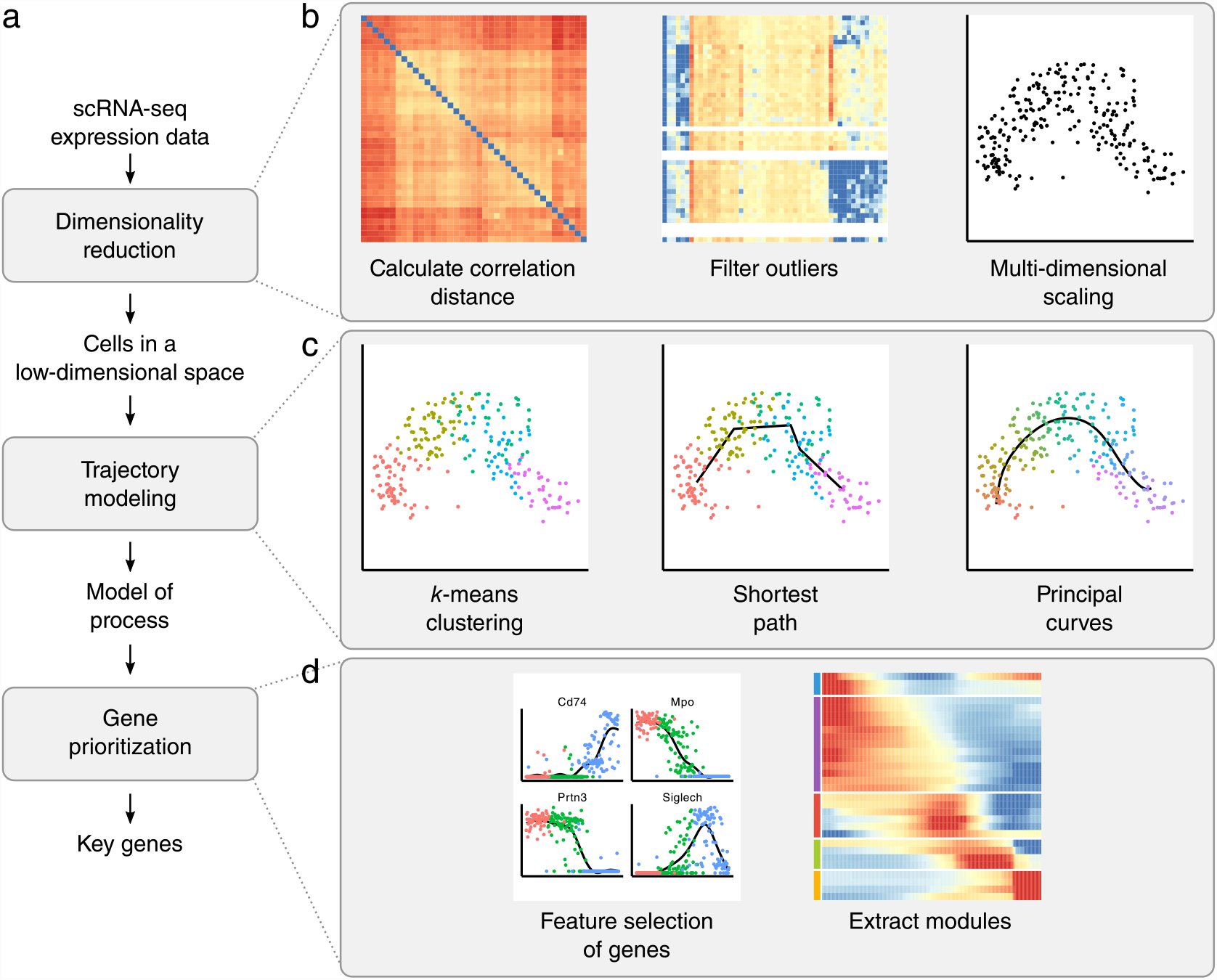
SCORPIUS infers a trajectory in three steps. 1) Dimensionality reduction involves calculating the correlation distance, optionally filtering out outliers, and performing multi-dimensional scaling. 2) Trajectory inference creates an initial path by calculating the shortest path through *k* cluster centres, and by iteratively fitting this path to the data using the principal curves algorithm. 3) During feature selection, a Random Forest is trained using the expression data to predict the ordering of cells as outputted by the principal curves. This Random Forest is used to select the most important genes in the dataset, cluster these, and check them for gene set enrichment.

In the second step, SCORPIUS reconstructs a trajectory through the data (Figure 1c). An initial trajectory is constructed by clustering the data with k-means clustering, and finding the shortest path through the cluster centers. This initial trajectory is subsequently refined in an iterative way using the principal curves algorithm [19]. The individual cells can then be ordered by projecting the n-dimensional points onto the trajectory. In the third and final step, SCORPIUS infers the degree to which a gene and its expression is involved in the dynamic process of interest (Figure 1c). This is achieved by ranking the genes according to their ability to predict the ordering of cells from the expression data, using the Random Forest algorithm [20]. The genes are then clustered into coherent modules, and visualized in order to improve the interpretability of the constructed model.

### 3.2 Extensive benchmarking shows SCORPIUS outperforms existing TI methods

At the time of introduction of the pioneering TI methods, the number of publicly available scRNA-seq datasets usable for investigating dynamic processes was severely limited, and thus the evaluations of these methods have been restricted to using only one or two datasets. The increasing number of publicly available scRNA-seq datasets now allows to perform a first extensive benchmarking experiment and thus quantitatively assess the performance of existing TI methods. We collected ten scRNA-seq datasets from five studies [8, 21, 3, 17, 12], representing several types of dynamic processes: cell differentiation, cell cycle and response upon external stimulus (See Table S1). For each of these datasets, labels regarding the state of cells in the dynamic process are available (e.g. using expression of known differentiation markers), which was used strictly only to evaluate a method, not to infer a model with. In this benchmark, SCORPIUS was compared with three state-of-the-art methods: Wanderlust [16], Monocle [17] and Waterfall [18]. A detailed overview of the characteristics of each approach can be found in Table S2.

Similarly to SCORPIUS, these alternative TI methods also first use a dimensionality reduction step and subsequently infer a trajectory in the reduced space (Supplementary Note). As shown in Figure 2a, we evaluated both these steps using two different metrics, respectively the accuracy and the consistency. The accuracy metric quantifies the performance of the dimensionality reduction step by measuring how accurate the cell labels are grouped together in the reduced space. To this end, the accuracy is calculated by predicting the label of each cell from its five nearest neighbors (5-NN), each time comparing the true cell label to the one predicted based on its five nearest neighbors. A good accuracy means that the reduced space has suffcient information to preserve cell state similarity. The consistency metric quantifies the performance of the trajectory inference step by comparing the predicted cell ordering to the known progression in the dynamic process. The consistency score is calculated by counting the number of consistent and inconsistent orderings for each cell in the trajectory with respect to the known progression, and is equal to the average percentage of consistent orderings per cell. Differences in scores due to stochastic components were removed by running each method on each dataset 100 times and averaging the scores.

**Figure 2:**
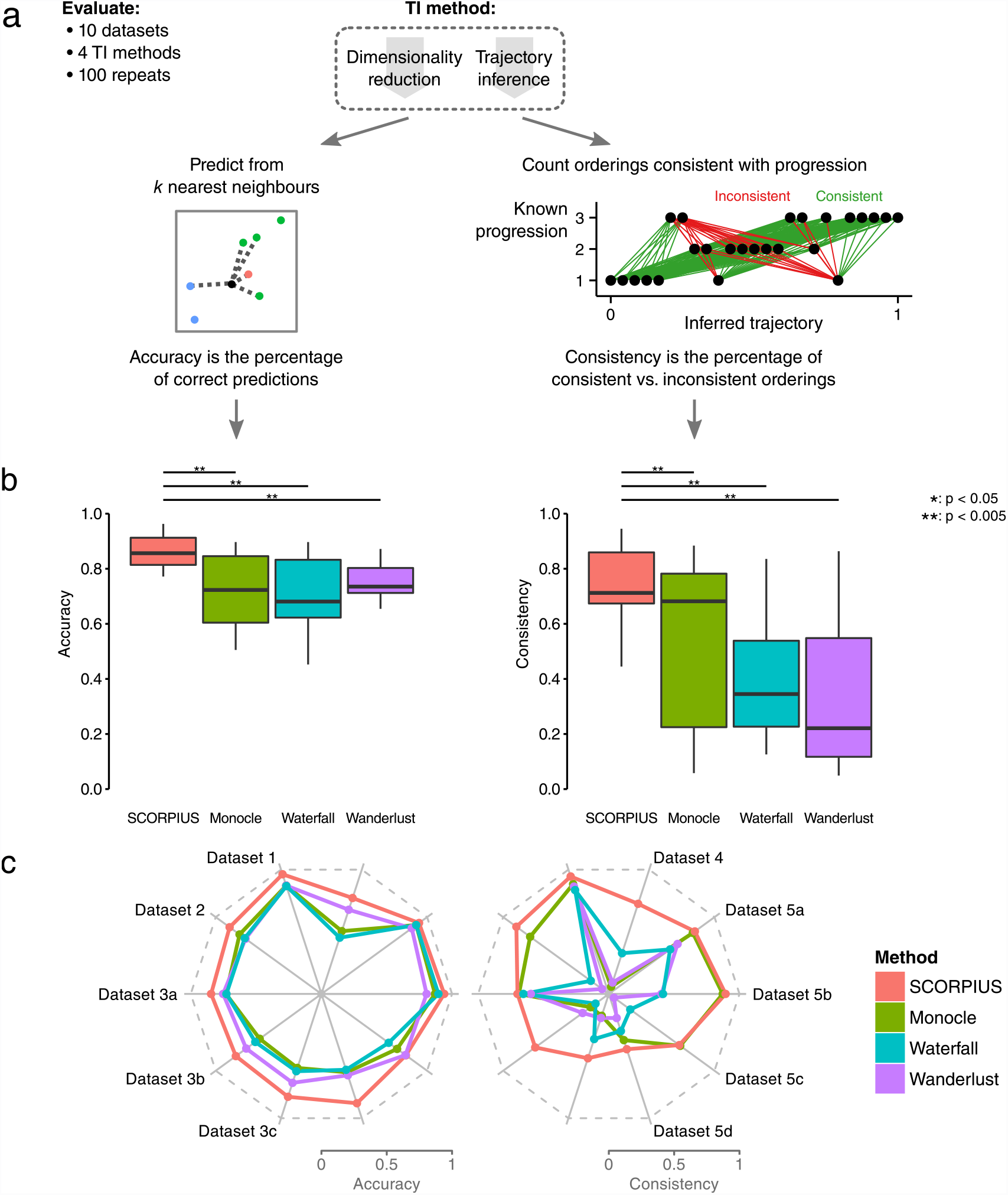
The workflow and results of the benchmarking experiments. a) For each of the TI methods, we evaluated two steps common to all approaches using 10 different datasets using the cross-validation accuracy (CVA) and consistent ordering score (COS). By predicting the progression stages of cells using a k-nearest-neighbors approach and calculating the accuracy of those predictions, the dimensionality reduction step is evaluated (left). A trajectory (right) is evaluated by counting which pairs of cells have an inferred pseudo-time consistent with the known progression. The consistency of a trajectory is the percentage of consistent orderings. b) SCORPIUS outperforms other state-of-the-art approaches, both in dimensionality reduction as well as the ordering of the cells (*: p-value < 0.05, **: p-value < 0.005). c) Accuracy and consistency scores for every method and dataset show that inferring accurate trajectories is more difficult for some methods.

SCORPIUS significantly outperforms other TI methods both in terms of accuracy and consistency (Figure 2b). It outperforms all other methods for each of the datasets (Figure 2c), except on dataset 5c, where Monocle achieved a higher consistency score in comparison to SCORPIUS. While the dimensionality reduction step of SCORPIUS generally performs well, its trajectory inference step performed worse on datasets 3c and 5d. Visual inspection of the inferred trajectories showed that dataset 3c contains a lot of heterogeneity between cells in the same time points, indicating the presence of subpopulations in the dataset, and that dataset 5d might contain a large systematic error, as each of the TI methods had ordered the ST-HSC and LT-HSC stages incorrectly. While all of the methods achieved a relatively high score on their dimensionality reduction steps, the performance of their trajectory inference steps is variable. For Monocle, this is to be expected, as calculating the longest connected path in a minimum spanning tree between cells is highly sensitive to noise [22]. Calculating the shortest-path distance from a starting node seemingly works well on some datasets and not on others, as the performances of Waterfall and Wanderlust are highly variable but very correlated.

### 3.3 SCORPIUS highlights different functional modules in dendritic cell development

In order to demonstrate the capability of SCORPIUS to generate testable hypothesis on real data, we applied the SCORPIUS algorithm on a recent scRNA-seq dataset of dendritic cell (DC) progenitors [8]. Although dendritic cells play a critical role in the activation of the adaptive immune system in vertebrates, several key regulatory mechanisms involved in this process are still disputed [23, 24]. DC progenitors are derived from hematopoietic stem cells in the bone marrow, and transition through a plethora of cellular states before becoming fully developed DCs (Figure 3a) [25, 26, 27, 28, 29, 30]. The dataset contains 57 Monocyte and Dendritic cell Progenitors (MDPs), 95 Common Dendritic cell Progenitors (CDPs) and 96 Pre-Dendritic Cells (PreDCs). SCORPIUS correctly orders the cells with regard to their differentiation status, as indicated by comparing the inferred trajectory with the known transition states (Figure 3b).

**Figure 3:**
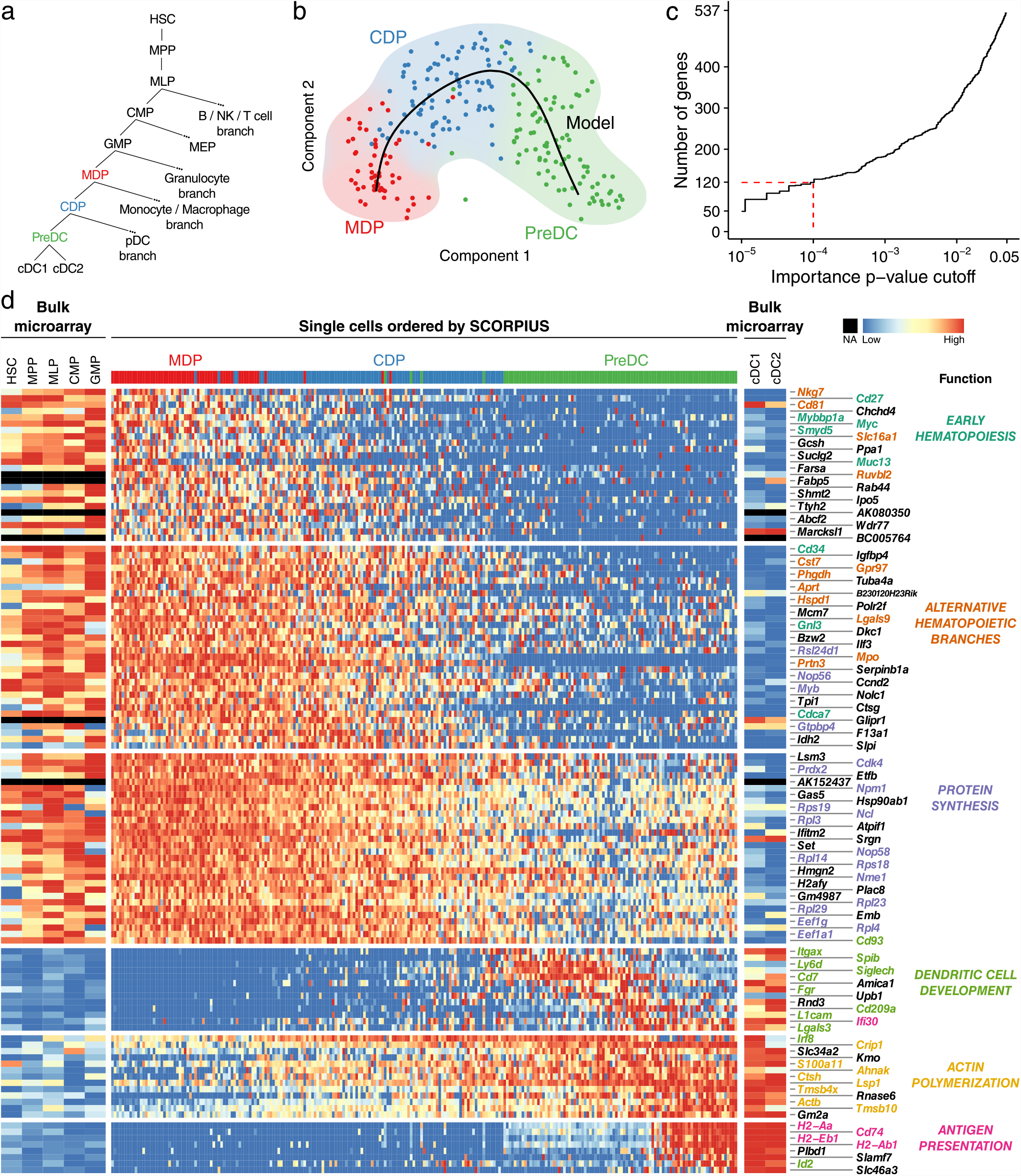
SCORPIUS sheds new, data-driven light on dendritic cell development. a) Dendritic cell precursors are derived from bone marrow stem cells and transition through many cell stages before finally becoming developed dendritic cells. b) SCORPIUS creates an accurate model for DC development from scRNA-seq data. c) The top 120 most important genes (p < 10-4) were retained for further investigation. d) These genes are clustered into six gene modules. Each module is responsible for different aspects of DC development. Bulk microarray expression for other stages of dendritic cell development is shown.

SCORPIUS then infers the degree to which a gene is involved in DC development, by using Random Forests [20] to predict the pseudotime ordering from the expression data and subsequently estimating the importance of each gene in this prediction. Empirical p-values were calculated by permutation testing (Figure 3c), and the most predictive genes (p < 10-4) were clustered into coherent gene modules (Figure 3d). The number of clusters was automatically determined with the Bayesian information criterion. We found that not only do the modules contain genes with very similar expression profiles, these genes also have very similar functions. In addition, further validation with bulk microarray expression data [31] shows a high similarity between the two data sources and gives insight into the expression of the selected genes for other cell types.

Modules 1 and 2 primarily contain genes that are involved in early hematopoiesis (e.g. *Cd27*, *Cd34*) or the development of a different hematopoietic lineage branch (e.g. NK cells: *Nkg7*; *myeloid*: *Mpo*, *Prtn3*; B cell: *Cd81*, *Gpr97*, *Hspd1*; T cell: *Nkg7*, *Cd81*, *Hspd1*, *Lgals9*). We found that while the expression of these genes was relatively high in the progenitor cells, it rapidly decreased during DC differentiation. A possible explanation could be that a suffcient level of proteins (which these genes transcribe for) has been reached, and that the mRNA expression is reduced in order to decrease the synthesis levels of the respective proteins.

Module 3 contains many genes related to protein synthesis (e.g. *Ncl*, *Cdk4*) which progressively decrease in expression during DC differentiation. As it is known that the protein synthesis rate gradually decreases during granulocyte and B-cell development [32], this module suggests that an analogous process exists during DC development.

Module 4 contains mostly genes that are already known to be involved in dendritic cell development (e.g. *Itgax*, *Cd209a* and *Lgals3*), confirming the ability of SCORPIUS to recover drivers of DC development in a purely data-driven way. It comes as no surprise that these genes are upregulated in the PreDC stage as these cells are preparing to become fully developed DCs.

The genes in module 5 are involved in actin polymerization (e.g., *Tmsb4x* and *Crip1*) and contains additionally one actin isoform (*Actb*). DCs rely on a filamentous actin cytoskeleton to capture antigens and facilitate locomotion [33, 34], the dynamics of which are determined by constant cycles of polymerization and depolymerization. While it is known actin polymerization plays a crucial role in the morphology, migratory behavior, and antigen internalization capacity of DCs, the upregulation of the genes in module 5 suggests that the synthesis of the proteins required for actin polymerization picks up during the CDP and PreDC stages. This shows the power of TI methods to exactly pinpoint the cellular states at which genes necessary for a particular cellular function get upregulated.

Module 6 contains mostly genes that are involved in antigen presentation (*Cd74*, *H2-Aa*, *H2-Ab1*, *H2-Eb1*), one of the major functions of DCs [35], as part of the major histocompatibility complex (MHC) class II. Whereas PreDCs have low MHC II expression on the cell surface, the high mRNA expression of these genes in late PreDCs indicates these cells are preparing to become developed DCs, to migrate and present antigen on their cell surface.

### 3.4 A decrease in protein synthesis rate during dendritic cell development is confirmed *in vivo*

The identification of module 3, containing genes related to protein synthesis, suggests that during DC development translation is decreased (Figure 4a), a novel hypothesis within the field of DC development. In order to verify whether translation indeed decreases during DC development, we quantified the protein synthesis rate of murine bone marrow cells *in vivo*. We intraperitoneally injected O-propargyl-puromycin (OP-Puro), an amino acid analogue, which enters ribosome acceptor sites and is incorporated into nascent polypeptide strands [36]. The subsequent fluorescent labeling of OP-Puro allows us to quantify the proteins synthesis rate on a single cell level using flow cytometry.

**Figure 4:**
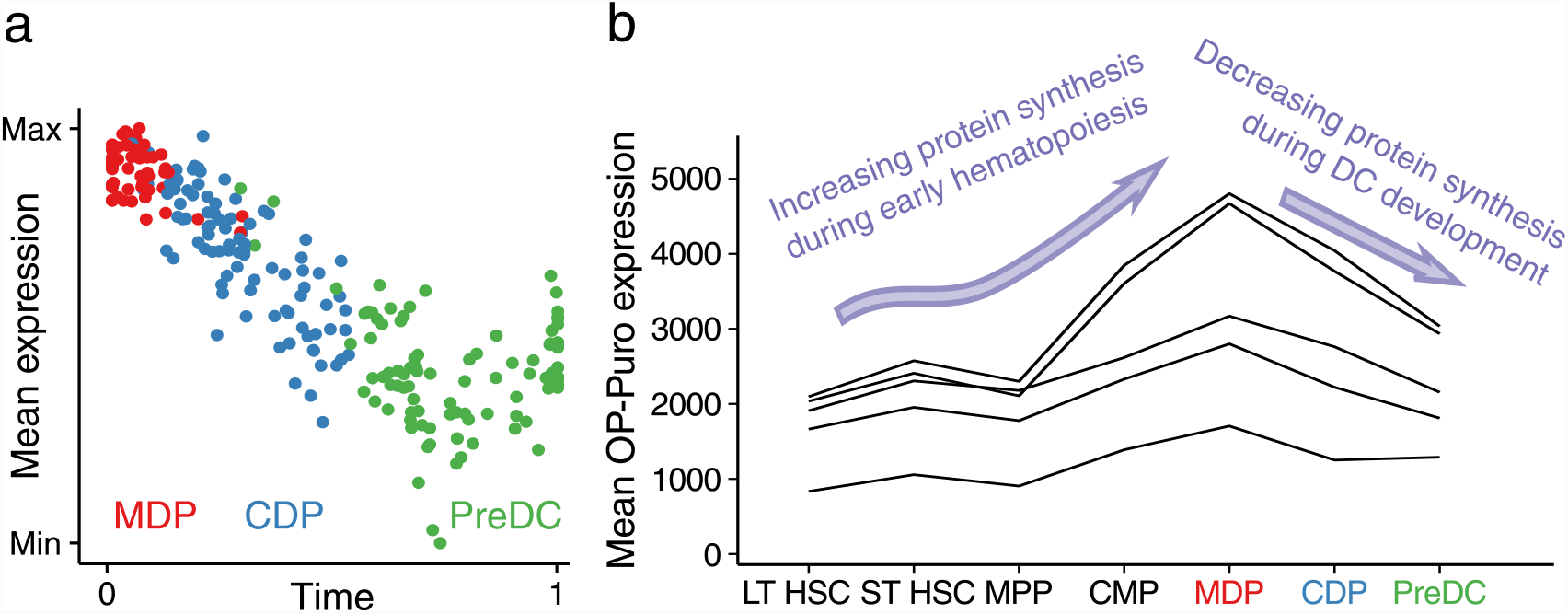
Protein synthesis is an integral part of DC development. a) Expression of genes involved in protein synthesis decreases during DC development. b) The protein synthesis rate of several stages of dendritic cell progenitors was measured with OP-Puro, showing MDPs have a high protein synthesis rate which is reduced throughout the differentiation process.

While the OP-Puro fluorescence intensities varied across the five individual mice, the relative fluorescence levels are very similar across replicates (Figure 4b). As described previously [32], OP-Puro incorporation is significantly lower in HSCs and multipotent progenitors (MPPs) than in common myeloid progenitors (CMPs). In line with the decreasing transcript expression levels of protein translation genes, the OP-Puro fluorescence levels and thus also protein production levels progressively decrease during DC development.

We summarize the results obtained by this work in the context of DC development by mapping the six coherent modules onto the DC lineage tree (Figure 5). We found that each of these modules corresponded to particular functions, which are up and down-regulated in different waves during DC development. While some of these functions have already been described as being important during DC differentiation [35, 33], such as the late upregulation of antigen presentation genes, our finding that translation related genes become downregulated during DC development was more unexpected. The functional activity of this finding was further confirmed through quantification of OP-Puro incorporation, thus demonstrating the capacity of SCORPIUS to construct high-quality models and generate testable hypotheses therefrom.

**Figure 5:**
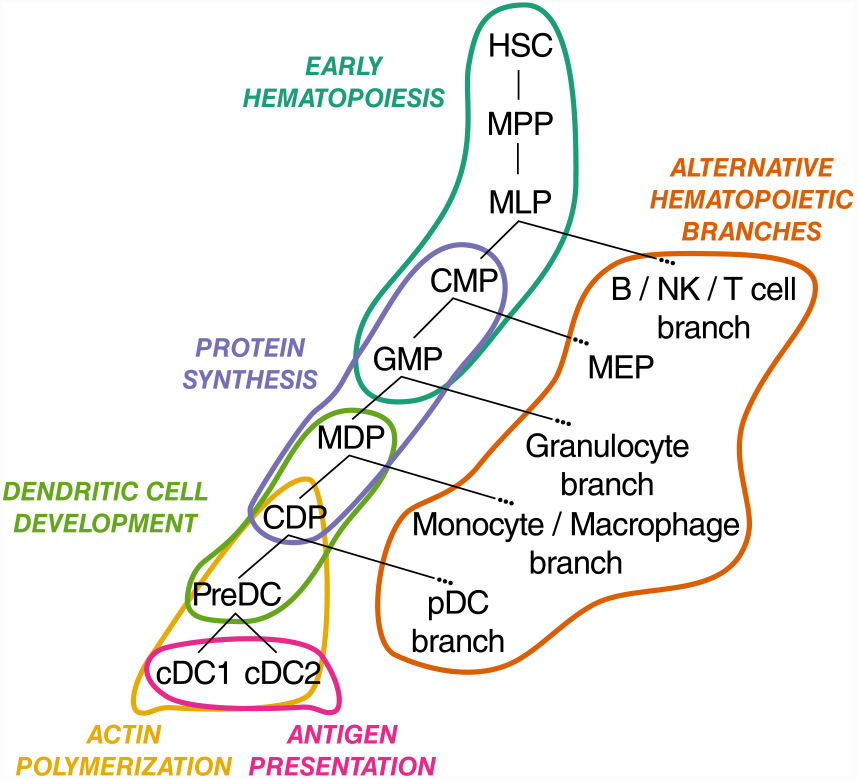
Gene dynamics during DC differentiation come in different functional waves. We mapped the activity of the processes related to each of the modules onto the dendritic cell lineage tree. Following the results from the OP-Puro experiment, we confirm that a dynamic protein synthesis rate is an integral part of DC development in CMPs, GMPs, MDPs and CDPs. The functional activity of other gene modules is in line with existing literature.

## 4 Discussion

New technological advances in the field of single-cell transcriptomics are revolutionizing the field of biotechnology, and classical clustering techniques are not designed to model dynamic processes that represent gradual changes in cell state. To better model such gradual changes, trajectory inference methods have recently been introduced to order single cells along a pseudo-temporal timeline implicitly present in the data. The resulting ordering can subsequently be used to infer novel dynamical properties of processes such as cellular development, differentiation and responses to stimuli. As most of the existing TI methods depend on prior biological knowledge, we propose SCORPIUS, a novel TI method that infers trajectories in a purely data driven way. In addition, as a large-scale quantitative evaluation of TI methods had hitherto been lacking, we developed a benchmarking strategy and found none of the existing TI methods performed well on all of the datasets consistently.

SCORPIUS was shown to be able to accurately infer trajectories from single-cell expression data on a wide range of datasets. While self-assessment can lead to an overestimation of the general performance of a method [37], we attempted to reduce the bias due to selective reporting of performance by benchmarking the SCORPIUS and other TI methods on ten different scRNA-seq datasets. We showed that SCORPIUS is able to consistently infer accurate models for different dynamic processes, and statistically outperforms existing TI methods.

We further validated the potential of SCORPIUS on real datasets by inferring an accurate model for the development of dendritic cells, a crucial type of antigen-presenting cells that bridge the innate and adaptive immune system. SCORPIUS identified well-known properties of DCs in a purely data-driven way, and using independent bulk microarray data we confirmed the up- and down-regulation of several modules during DC development. Through the observation of a decrease in mRNA expression of genes involved in translation, we hypothesized that protein synthesis levels progressively decrease throughout development, a new finding regarding DC development. We quantified the protein synthesis rates of various DC progenitors and confirmed high translational activity of MDPs which decreases steadily. Translation of proteins is a highly regulated process throughout the development of both hematopoietic and non-hematopoietic cells [38, 32] cells display low rates of protein synthesis, only increasing upon generation of rapidly cycling cell types such as common myeloid progenitor cells (CMPs) and MDPs to support their higher proliferative capacity [32]. It is believed that this tight regulation of protein synthesis in stem cells, through the PERK-eIF2a axis, is important for protection against stress associated with protein folding and to preserve stem cell longevity [38, 39]. Naik et al. reported reduced proliferative capacity of CD11c+ PreDCs compared to CD11c-MDPs and CDPs [30]. Thus, our observations of reduced translational machinery in PreDCs can be interpreted along similar lines, suggesting that cells such as PreDCs reduce the translational capacity according to their needs.

An apparent shortcoming of the inferred trajectories is the assumption that the dynamic process of interest is linear. The results obtained by this study also indicate the necessity of being able to infer branching or even more generalized models. While inferring a linear model is relatively simple in terms of the model complexity, inferring a branching or generalized model is considerably more complex, and thus requires a much greater number of cells. Up until recently it was only possible to sequence the transcriptomic profiles of hundreds of single cells, but new developments in single cell RNA sequencing allow thousands or more single cells to be profiled and thus also the construction of more complex models. This has resulted in the introduction of novel branching TI methods [40, 41, 42]. A large-scale quantitative benchmarking of these methods is greatly warranted, but will require the development of novel performance metrics and collection of scRNA-seq datasets investigating branching trajectories.

A second challenge highlighted by this study is the effect of simultaneous dynamic processes, for example maturation and cell cycle. While one approach could be to remove effects from cell cycle by correcting the expression levels [21], simultaneous dynamic processes are likely to interact with each other, and thus removing the effects of one of those processes can be detrimental to downstream analyses. A better approach would be to attempt to separate variation of expression into several dynamic processes, but this again requires larger datasets.

Index sorting is an exciting new development when constructing computational models from single cell data. By sorting single cells into individual wells using a flow cytometer, index sorting allows to obtain a transcriptomic and proteomic profile of each cell. This is inter alia extremely useful when investigating dynamic processes, as this will improve accuracy of the model as increases in protein levels should be preceded by increases in mRNA levels. While the selection of cells in cytometry is defined by gating structures, index sorting also allows to capture cells throughout the whole spectrum of the dynamic process of interest, thereby removing gates as a potential source of bias in the experiment.

In summary, this work introduces a novel approach for inferring computational models of linear dynamic processes in an accurate and data-driven approach. Careful design of the methodology and the quantitative evaluation play a crucial role in reducing bias in the models that are inferred. In doing so, this work enables de novo investigation and characterization of dynamic processes and lays the foundation for objective benchmarking of future trajectory inference methods.

## Methods

### Code availability

All code used in this study is made publicly available allowing the replication of analyses and enabling easier and more robust benchmarking strategies. An open source implementation of SCORPIUS is available on GitHub: github.com/rcannood/SCORPIUS.

### Benchmark data sets

We collected 10 scRNA-seq datasets representing several types of dynamic processes: cell differentiation, cell cycle and response upon external stimulus. An scRNA-seq dataset had to contain cells at different stages as part of a dynamic process for which the labels were experimentally determined, and at least 50 cells had to be present per progression stage. For each of the datasets, we downloaded expression data and extracted progression labels from the respective accession codes listed in Table S1. Aside from log-transforming the expression values, no further preprocessing of the expression data was performed.

### Dimensionality reduction

Define ℂ as the collection of all cells. The distance between any two cells is defined as:

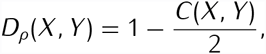

where *C*(*X, Y*) is the Spearman’s rank correlation for tied ranks (Zar 2005). We define the *outlierness* of a cell as the mean distance to it and its 10 nearest neighbors:

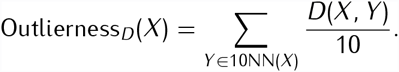

We assume the outlierness of all cells to be normally distributed. Thus, we iteratively remove cells with maximal outlierness and fit a normal distribution to the remaining values using the *fitdistrplus* R package. Finally, we ultimately retained those cells at which the log likelihood of the fit is maximal. In order to reduce a dataset to an n-dimensional space S, we perform classical Torgerson multi-dimensional scaling:

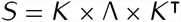

where ⋀ is a vector containing the *n* largest eigenvalues of the double-centered *D*_*ρ*_, and *K* is a matrix containing the corresponding eigenvectors. All results in this study were produced with *n* = 3.

### Trajectory inference

First, the cells are clustered into *k* clusters with *k*-means clustering within the reduced space *S*. All results in this study were produced with *k* = 4. Next, an initial rough estimate of the trajectory is searched for by linking cell clusters through their shortest path using a custom distance function. The distance function takes into account the distance between two cluster centers, as well as the density of cells between the two cluster centers. It is defined as:

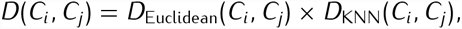

with *D*_Euclidean_ defined as the Euclidean distance between clusters *C*_*i*_ and *C*_*j*_, and *D*_KNN_ defined as the mean distance of evenly spread points between *C*_*i*_ and *C*_*j*_ and their respective 10 nearest neighbors (defined earlier as the outlierness):

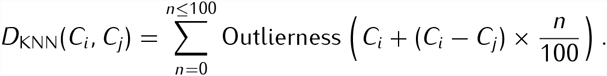

Finally, SCORPIUS further optimizes this initial path using the Hastie and Stuetzle principal curves algorithm [43] (implemented in the *princurve* R package). This algorithm iteratively smoothens the trajectory until convergence. Each iteration, the cells are projected onto a given curve, and a new curve is constructed by locally averaging the projected cells.

### Feature selection

We used the Random Forest [20] algorithm to assess the importance of a gene with respect to the inferred trajectory. By using a random forest to predict the pseudotime of a cell from its expression data, the importance of a gene’s expression with respect to the prediction made can be calculated.

More specifically, a random forest consists of many decision trees in which non-leaf nodes represent decision splits based on one of the genes and leaf-nodes contain predictions for the orderings. The importance of a gene is then the mean decrease in mean squared error (MSE) each time that gene is used to create a split. The genes are ordered by importance.

Gaussian mixture models were used to cluster the expression of the top genes into modules. These modules were initialised with hierarchical clustering, and were optimised with the Bayesian information criterion (BIC). The implementation of this approach is provided by the *mclust* R package [44].

### Evaluation metrics

The dimensionality reduction and trajectory inference steps of a method were evaluated using the *cross-validation accuracy* (CVA) and the *consistent ordering score* (COS) metrics, respectively. Define *E*_*I*_ as the experimentally observed progression of a cell *I ∈* ℂ, and *O*_*I*_ as its ordering along an inferred trajectory.

The accuracy score is calculated by predicting the progression label of each cell from its 5 nearest neighbours and calculating the percentage of correct predictions:

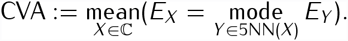

The consistency score was defined as the percentage of pairwise orderings within the trajectory which is consistent with the known progression. Since the direction of the trajectory is not inferred, the absolute value of the consistency score is used.

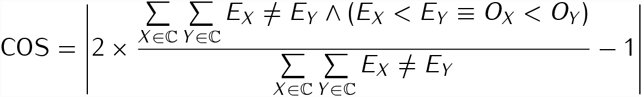

### Performance comparison

We compared SCORPIUS with three other TI methods: Wanderlust, Monocle, and Waterfall. The overall characteristics of these methods are listed in Table S2. For each of the methods, the default parameters were used (Wanderlust: num_landmarks = 20, num_graphs = 100, k = 30, l = 8; Monocle: num_genes = 1000, num_paths = 1; Waterfall: k = 5). In order to be able to compare the scores between methods, the same outlier filtering was used for each of the TI methods. Each TI method was executed 100 times on each dataset, and the mean CVA and COS was calculated. To determine the significance values of differences in performance, we performed a one-sided paired Wilcoxon rank test.

### Measurement of protein synthesis

O-Propargyl Puromycin (Jena Bioscience - NU-931-5) was dissolved in DMSO, further diluted in PBS (10 mg mL^*−*1^) and injected intraperitoneally (50mg/kg mouse weight). 1 hour after injection mice were euthanized by cervical dislocation and hind bones were collected. Bone marrow cells were obtained by crushing of bones with pestle and mortar and subsequent lysis of red blood cells. The remaining cells were filtered through a 70 µm mesh and resuspended in a Ca^2+^ and Mg^2+^ free phosphate buffered solution (PBS; Gibco). Viable cell numbers were assessed with a FACS Verse (BD Biosciences).

7 × 10^6^ cells were stained with mixtures of antibodies directed against cell surface markers. Each staining lasted approximately 30 min and was performed on ice protected from direct light. Monoclonal antibodies labeled with fluorochromes or biotin recognizing following surface markers were used: CD3 (145-2C11; Tonbo), TCRb (H57-597; BD Pharmingen), CD4 (RM4-5; eBioscience), CD8a (53-6.7; BD Pharmingen), CD19 (1D3; Tonbo), CD45R (RA3-6B2; BD-Pharmingen), TER119 (TER119; eBioscience), Ly-6G (1A8; BD-Pharmingen), NK1.1 (PK136; eBioscience), F4/80 (BM8; eBioscience), CD11c (N418; eBioscience), MHCII (M5/114.15.2; eBioscience), CD135 (A2F10; eBioscience), CD172a (P84; eBioscience), CD45 (30-F11; eBioscience), SiglecH (eBio440c; eBioscience), Ly-6C (HK1.4; eBioscience), CD115 (AFS98; eBioscience), CD117 (2B8; eBioscience), CD127 (SB/199; BD-Pharmingen), Ly-6A/E (D7; eBioscience), CD34(RAM34; eBioscience), CD11b (M1/70; BD Pharmingen). Viable cells were discriminated by the use of the fixable viability dye eFluor506 or eFluor786 (eBioscience).

Next, cells were fixed and permeabilized using the FoxP3 Fixation/Permeabilization kit (eBioscience, 00-5521-00). For OP-Puro labeling, Azide-AF647 is chemically linked to OP-Puro is through a copper-catalyzed azide–alkyne cyloaddition. In short, 2.5 µM azide-AF647 (Invitrogen, A10277) is dissolved in the Click-iT Cell Reaction Buffer (Invitrogen, C10269) containing 400 µM CuSO_4_. Immediately after preparation, cells are incubated with this mixture on room temperature. After 10 min incubation, the reaction is quenched by addition of PBS supplemented with 5% heat-inactivated fetal calf serum (FCS; Sigma) and 5mM EDTA (Lonza; 51234). Cells are washed twice to remove unbound azide-AF647. A Fortessa X20 (BD Biosciences) was used for data acquisition and data was analyzed using FlowJo 10 (LLC).

## Author Contributions

Conceptualization, R.C., W.S., and Y.S.; Methodology, R.C., W.S., and Y.S.; Software, R.C.; Investigation, R.C., S.T., and D.S.; Resources, S.J., M.G., and B.L.; Writing – Original Draft, R.C.; Writing – Review & Editing, R.C. W.S, Y.S, S.T., D.S., S.J., M.G., B.L., and K.D.P.; Supervision, Y.S., B.L., and K.D.P.

## Acknowledgements

This work is supported by Fund for Scientific Research FWO Flanders (R.C. and W.S.). Y.S. is an ISAC Marylou Ingram scholar.

## Supplementary Information

**Table S1:** Overview of the scRNA-seq datasets used in this study. Datasets had to contain cells at different stages as part of a dynamic process for which the labels were experimentally determined, and each stage had to contain at least 50 cells. We used 10 datasets originating from 5 different studies, for which the progression labels were determined through cell sorting or by sampling cells at different time points. Three different dynamic processes are investigated in these datasets: differentiation, cell cycle and response upon external stimulation.

| # | Ref. | Accession | Dynamic process | Cells | Progression stages | Source | #cells |
| --- | --- | --- | --- | --- | --- | --- | --- |
| 1 | Schlitzer et al. 2015 | GSE60783 | Differentiation | Pro-DCs | MDP ▷ CDP ▷ PreDC | FACS | 242 |
| 2 | Buettner et al. 2015 | E-MTAB-2512 | Cell cycle | mESCs | G1 ▷ S ▷ G2/M | FACS | 266 |
| 3a-c | Shalek et al. 2014 | GSE48968 | Response | DCs | 1h ▷ 2h ▷ 4h ▷ 6h | Time | 897 |
| 4 | Trapnell et al. 2014 | GSE52529 | Response | HSMMs | 0h ▷ 24h ▷ 48h ▷ 72h | Time | 285 |
| 5a-d | Kowalczyk et al. 2015 | GSE59114 | Differentiation | HSCs | LT-HSC ▷ ST-HSC ▷ MPP | FACS | 1535 |

**Table S2:** The strengths and weaknesses of each of the TI methods. Biological validation in study: + yes, – no. Quantitative evaluation in study: + yes, – no. Consistent performance in benchmarks: + yes, – no. Uses no prior knowledge: + yes, – no. Easy to use: + high quality code and documentation was provided, – code and/or documentation was not provided. Scalability of the method with respect to the number of cells or genes was determined by executing each method on an increasing number of cells until the execution time exceeded 10 seconds: + >10.000, ± <10.000, – <1.000.

|  | Wanderlust | Monocle | Waterfall | SCORPIUS |
| --- | --- | --- | --- | --- |
| Biological validation in study | + | + | + | + |
| Quantitative evaluation in study | - | - | - | + |
| Consistent performance | - | - | - | + |
| Uses no prior knowledge | - | - | + | + |
| Easy to use | - | + | - | + |
| # cell scalability | - | - | ± | ± |
| # gene scalability | + | ± | + | + |

